# Ultra-fine temporal resolution in auditory processing is preserved in aged mice without peripheral hearing loss

**DOI:** 10.1101/363994

**Authors:** Lasse Osterhagen, K. Jannis Hildebrandt

## Abstract

Age-related hearing loss (presbycusis) is caused by damage to the periphery as well as deterioration of central auditory processing. Gap detection is a paradigm to study age-related temporal processing deficits, which is assumed to be determined primarily by the latter. However, peripheral hearing loss is a strong confounding factor when using gap detection to measure temporal processing. In this study, we used mice from the CAST line, which is known to maintain excellent peripheral hearing, to rule out any contribution of peripheral hearing loss to gap detection performance. We employed an operant Go/No-go paradigm to obtain psychometric functions of gap in noise (GIN) detection at young and middle age. Besides, we measured auditory brainstem responses (ABR) and multiunit recordings in the auditory cortex (AC) in order to disentangle the processing stages of gap detection. We found detection thresholds around 0.6 ms in all measurement modalities. Detection thresholds did not increase with age. In the ABR, GIN stimuli are coded as onset responses to the noise that follows the gap, strikingly similar to the ABR of noise bursts in silence (NBIS). The simplicity of the neural representation of the gap together with the preservation of detection threshold in aged CAST mice suggests that GIN detection in the mouse is primarily determined by peripheral, not central processing.

**Abbreviaions:**

GIN: gap in noise
ABR: auditory brainstem response
AC: auditory cortex
NBIS: noise burst in silence
IIN: inhibitory interneuron

## 1 Introduction

Presbycusis is the progressive loss of hearing abilities due to aging. Two mechanisms are presumed to underlie age-related hearing impairment: damage to the peripheral sensory organ, which results in partial deafferentation, and deterioration of processing at later stages along the auditory pathway. Identifying the contribution of central processing deficits to presbycusis poses difficulties, because many elderly have concomitant peripheral hearing loss and/or more general cognitive impairments that may corrupt their performance during audiometry (Humes et al. 2012).

The detection of short gaps within sound is a frequently used paradigm to study age-related impairment of temporal auditory processing in humans (e. g. B. C. J. Moore, Peters, and Glasberg 1992; Poth, Boettcher, Mills, and Dubno 2001; Snell 1997; Strouse, Ashmead, Ohde, and Grantham 1998), as well as in animals e. g. (Barsz, Ison, Snell, and Walton 2002; Boettcher, Mills, Swerdloff, and Holley 1996; Hamann, Gleich, Klump, Kittel, and Strutz 2004). There is some agreement that gap detection is a paradigm for non-peripheral hearing loss and that the deficits in the elderly are caused by the deterioration of central auditory processing. However, because of the difficulty in disentangling peripheral from central processing deficits, studies that employ the gap detection paradigm are not easy to interpret.

Behavioral gap detection performance strongly depends on the frequency content of the sound in which gaps are inserted: the higher the frequency, the shorter the gaps that can be detected (Fitzgibbons 1983; Green and Forrest 1989; Radziwon et al. 2009; Shailer and B. C. J. Moore 1983). Since the sensitivity for high frequencies declines with age (Agrawal, Platz, and Niparko 2008), high-frequency hearing loss is an important confounding factor when using gap detection performance as an indicator for central processing deficits. Though various attempts have been made to minimize the influence of this confounding factor (see discussion), it would be desirable to have a way to entirely exclude the possibility that peripheral hearing loss has an influence on gap detection performance.

Animal research provides fine-grained access points for the measurement and manipulation of auditory processing along the auditory pathway that are unavailable in research on human subjects. In particular, the importance of mice as an animal model in auditory research is increasing, because they allow various approaches for manipulation (e. g. genetic and optogenetic approaches) that are not available in other animal models. Mice from the B6.CAST-Cdh23 line are among those with the best hearing. Furthermore, they maintain excellent peripheral hearing throughout their life span (Zheng, Johnson, and Erway 1999). This makes them suitable to study central presbycusis with the gap detection paradigm, without the risk of contaminating the results by peripheral hearing loss.

In this study, we combine behavioral psychophysics by means of operant conditioning, ABR, and multiunit recordings in the AC in order to disentangle processing stages of gap detection. Furthermore, we investigate the effect of aging on gap detection. We use B6.CAST-Cdh23 mice to rule out any possibility that peripheral hearing loss has an influence on the results. Psychometric gap detection functions are measured behaviorally with a group of these mice at two ages. If we find the performance to be decreased at the older age, it will be most likely due to the deterioration of central processing.

## 2 Methods

### 2.1 Subjects

The animals we used in all experiments were male mice with a B6.CAST-Cdh23 background (Jackson Laboratory, Bar Harbor, ME). For these animals, it is known that they do not develop age-related peripheral hearing loss. Behavioral measurements took place when the mice were 25 weeks old and were repeated with the same animals at the age of 44 weeks. ABR measurements were obtained in the same group in a terminal experiment when the mice were 62 weeks old. Cortical recordings were obtained from animals between the ages of 20 and 25 weeks.

All animal experiments were done in accordance with the animal welfare regulations of Lower Saxony and done with the approval from the local authorities (State Office for Consumer Protection and Food Safety / LAVES), permission number 33.9-43502-04-13/1271.

### 2.2 Behavioral procedure

In order to obtain psychometric curves of behavioral GIN detection, we used the method of constant stimuli embedded in a Go/No-go paradigm with operant conditioning.

Experiment control and stimulus generation were performed by our PsychDetect software framework for MATLAB (The MathWorks, Natick, MA) running on a Microsoft Windows PC. Within PsychDetect, sound stimulation was generated digitally at a sample rate of 192 kHz. We used a 192 kHz, 24 bit USB sound device (Fireface UC, RME, Haimhausen, Germany) for D/A conversion. The sound was amplified by a PM7004 amplifier (Marantz, Kawasaki, Japan) and played back through a Vifa/Peerless XT-300 K4 (Tymphany, Sausalito, CA) loudspeaker.

Band-passed Gaussian white noise (flat frequency spectrum, range 2–64 kHz) calibrated to 50 dB SPL was continuously played throughout the session. Gap stimuli were inserted into the noise stream by setting sound samples to zero and applying a 0.3 ms cosine fading gate at the beginning and the end of the gap. Gap onset and offset were defined where the cosine gate attenuated the noise amplitude by one-half. Subjects’ detection performance was evaluated at six fixed gap length: 0.3, 0.4, 0.6, 1, 2, and 4 ms.

Training and testing of the subjects took place in a doughnut-shaped cage with a diameter of 30 cm. A 2.5 by 5 cm rectangular elevated podium was installed at one side of the cage, opposite to a feeding dish, that can be filled by an external feeder apparatus. The podium was located 45 cm below the loudspeaker. Subjects initialized trials by ascending the podium which was detected by a light barrier. After a waiting interval (one of 1.25, 2.25, 3.25, 4.25, and 5.25 s), a gap stimulus was inserted into the noise and the subjects had to leave the podium within 1 s in order to let the feeder release a food pellet into the feeding dish. If the subject left the platform during the waiting interval, the trial was canceled. When the subject again ascended the platform, the same trial was reinitialized.

An experimental session was constructed by selecting four gap lengths, two of the three shorter gap lengths and two of the three longer ones, out of the set of fixed gap lengths. A complete experiment consisted of all possible sessions that can be obtained according to this rule. (There are nine: three over two times three over two.) Within a session, each gap length was presented 15 times with equally distributed preceding waiting intervals. In addition, 25 sham trials were included. These are trials in which no stimulus was presented and a descent from the podium during the 1 s reaction time window was counted as a false alarm. Thus, a session consisted in total of 85 trials, which were randomly arranged. A session typically lasted between 25 and 50 minutes, depending on the subject’s motivation.

For each session, we computed hit rates for every gap length that was presented during that session and the session’s false alarm rate. Using signal detection theory, we combined these measures into *d′* values. Sessions were excluded from further analysis, if the subject did not reach a minimum *d′* of one or the false alarm rate exceeded 30 percent.

Four subjects of the same litter were trained prior to the first experiment by means of shaping, which required two weeks with two training sessions per day. The measurement of a psychometric function took five days with two sessions per day. The mice were 25 weeks old when the first measurement took place. To assess the influence of aging on gap detection performance, we repeated the measurement when the mice were 44 weeks old.

### 2.3 ABR procedure

Animals were anesthetized with Ketamine (50 mg/kg) and Medetomidine (0.415 mg/kg). Needle electrodes were placed subcutaneous with the recording electrode at the neck and the reference electrode at the vertex. The electrode signal was bandpass filtered (300 Hz–30 kHz) and amplified by an ISO-80 Bio-amplifier (World Precision Instruments, Sarasota, FL), before A/D-converted through a Fireface UC 24 bit sound device at a sample rate of 96 kHz. The same device was used for stimulus delivery. Stimuli were created by a custom MATLAB application.

The sound was delivered to the animals binaurally directly into the ear canal through horns that were attached to Vifa/Peerless XT-300 K4 loudspeakers. (For a detailed description see Beutelmann, Laumen, Tollin, and Klump 2015). The system was calibrated by means of small microphones (Knowles FG-23329, Itasca, IL) that were inserted into replicas of mice ear canals.

Just as in the behavioral procedure, we used continuous Gaussian white noise throughout the recording of ABRs for GIN stimuli. However, due to the limited frequency bandwidth of the calibration microphones, we bandpass filtered the noise between 2 and 46 kHz. Gaps were inserted into the noise stream as described before. The time between consecutive gap stimuli varied randomly between 50 and 150 ms. The sequence of stimulus repetitions was random. In a pilot study, we determined the best signal-to-noise ratio for GIN ABRs to be when the noise was played at 60 dB SPL. That means it requires the least number of stimulus repetitions to create average ABR curves. Therefore, the results presented here are typically derived from gaps in 60 dB SPL noise. Only for the animals that took part in the behavioral experiment, we also conducted ABR measurements for gaps in 50 dB SPL noise and and their ABR data presented here are from those recordings.

Besides GIN stimuli, we played 10 ms noise bursts in silence (NBIS), which were created likewise from Gaussian white noise (2–44 kHz bandwidth) and a fading gate of 0.3 ms, to allow the direct comparison of the ABRs from the gaps and the noise bursts.

Thresholds for artifact rejection (heart or breathing muscle potentials) were defined manually on the basis of test recordings prior to the ABR recordings. For every stimulus, at least 400 artifact-free repetitions were collected to create average ABR waveforms.

### 2.4 Cortical recordings

Neural responses from primary auditory cortex (A1) were obtained from four animals, aged between 4 and 8 months. A custom made array consisting of eight twisted-wire tetrodes (platinum-iridium, 37 μm diameter) attached to a miniature microdrive (Axona, London, UK) was implanted into just above right A1. Placement of the electrode tips was done according to stereotaxic coordinates obtained from previous experiments. Electrodes could be moved along a medial-lateral path tangential to the cortical surface. Recordings were made mostly from middle layers (IV). Placement of electrodes was confirmed by histology after finishing the experiments. Recordings were made using 32-channel digitizing head stage connected to an acquisition system (both OpenEphys, www.open-ephys.org). Continuous raw voltage traces were recorded to a PC and saved for offline-analysis. Recordings followed this schedule: on recording days, electrodes were moved ≈ 65 μm forward and allowed for 1–2 hours to settle. After this period, animals were connected to the recording system and placed in the acoustic booth and stimuli were presented with the stimuli. Animals were placed on a horizontal running wheel made out of wire-mesh in order to be acoustically transparent. Animals were free to run on the running wheel during the recording. Recordings in each animal were obtained from 6–10 positions in A1 along the medial-lateral axis.

#### 2.4.1 Stimulation

Stimuli consisted of 2 s noise bursts, in which a gap of varying duration was inserted, starting 1 s after noise onset. Gap durations of 0, 0.5, 1, 2, 4, 8, 16, 32 and 64 ms were used in all experiments. Noise stimuli were created offline by filtering broadband Gaussian white noise between 4 and 64 kHz. Stimulation was controlled by custom MATLAB software. The sound equipment was the same as used in the behavioral experiments. The speaker was mounted 1 m above the animal’s head and noise bursts were calibrated to 60 dB SPL at the animal’s ear position. Individual noise bursts were separated by 1.5 s of silence. All gap durations were presented in random order, each gap length was repeated 30 times.

#### 2.4.2 Analysis

Neural recordings were saved at 30 kHz sampling frequency for offline spike-sorting. Only well-separated single unit clusters were used for consecutive analysis. For analysis of neural gap thresholds, only units that showed significant responses to at least one gap duration were used for further analysis (261/743 single units from four animals). In order to obtain peri-time histograms (PSTHs), spike times were binned into 10 ms windows, starting at stimulus onset corrected for latency. A response to a gap was classified as significant if the spike count in any window between gap onset and gap onset + 100 ms was above a significance threshold. The time window was chosen in order to include both responses to the gap on- and offset for all stimuli (longest gap duration was 64 ms). The significance threshold was determined as the mean plus three standard deviations of the spike counts in reference windows during the noise stimulus before the gap, excluding the onset of the noise (400 ms after noise onset until gap onset). Since 10 windows were checked for significant threshold crossings against the distribution of reference windows, an additional factor of 1.23 based on correction for multiple comparisons was multiplied with the threshold. In addition to the algorithmic approach, all single responses of the units were visually inspected.

## 3 Results

### 3.1 Behavioral performance in young animals

We aimed to determine gap detection thresholds by an operant condition paradigm. To this end, we trained animals to indicate the presence of a gap in continuously presented noise by descending from a small platform in order to receive a food reward.

We measured behavioral GIN detection in four subjects for the first time, when the animals were 25 weeks old. According to our criteria, we had to exclude four sessions (two sessions from two subjects), because they exceeded our maximum false alarm rate. For each subject, median d′ values across sessions were computed for the six fixed gap durations (Figure 1). If we define the detection threshold at *d′* = 1, two subjects had a threshold below 0.6 ms gap duration (*d′* = 1.28 and 1.09 at 0.6 ms), while the other two subjects had a threshold slightly higher than 0.6 ms (*d′* = 0.91 and 0.84 at 0.6 ms).

**Figure 1:**
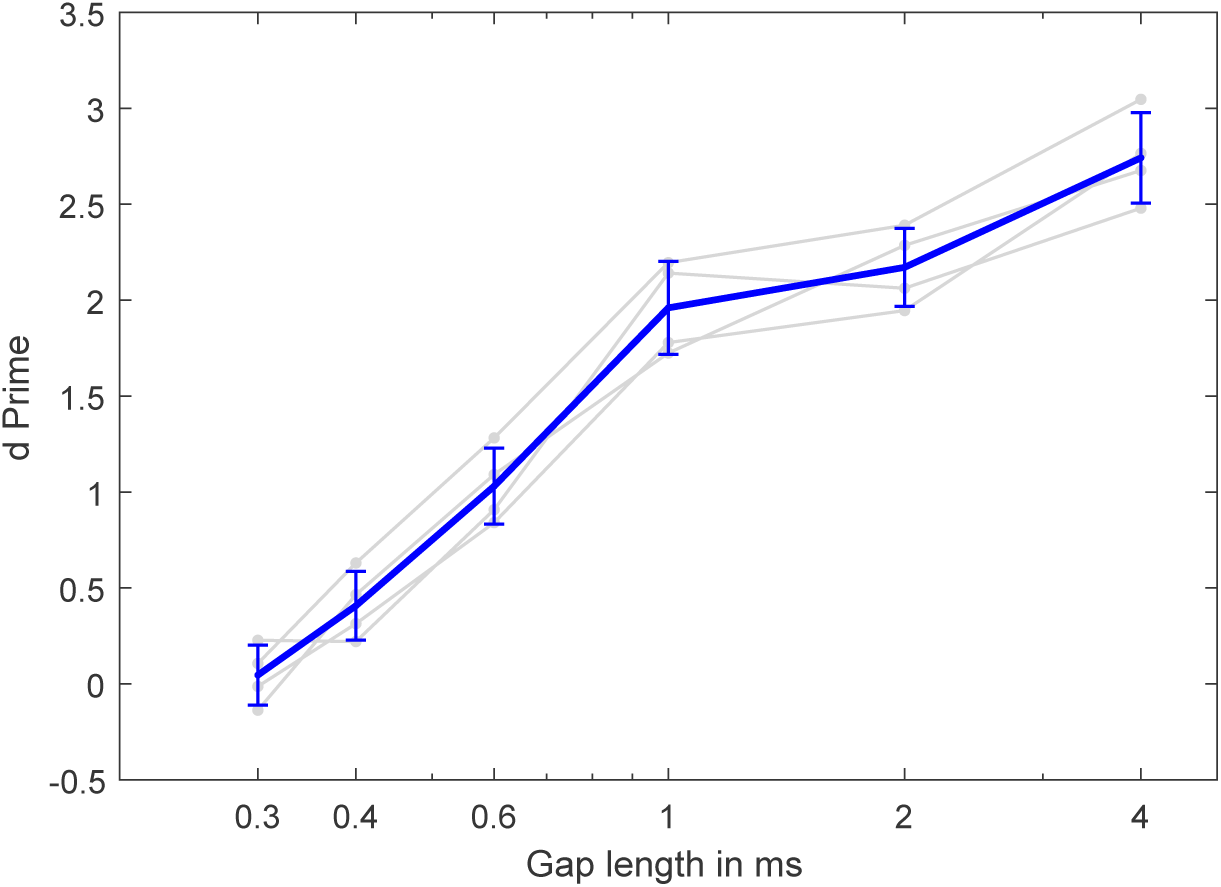
Psychometric functions for behavioral detection of GIN stimuli. The measurement took place when the mice (N=4) were 25 weeks old. *d′* sensitivity values are shown as a function of gap duration. Grey lines show data of individual subjects with the dots representing their median across sessions. The blue line displays mean *d′* (mean of subjects’ session medians) with error bars representing ± one SEM.

### 3.2 Cortical recordings

Since we found exceptionally low behavioral gap detection thresholds, we next sought to investigate neural basis of the detection. At which level can we find gap thresholds as low as we observed in the behavior? Since several studies highlight the importance of auditory cortex for gap detection, we started at this level of the auditory pathway to look for a possible representation of gaps in noise. We recorded from the primary auditory cortex (A1) of four awake, freely moving mice while presenting broadband noise with gaps varying from 0.5 to 64 ms. 35.1 % (261 out of 743) of units showed a significant response to at least one of the gaps. For these units, we determined the shortest gap to which the unit would respond. Figure 2 show raster plots for two example units, one with a high neural gap detection threshold (**A**, 32 ms) and one with a low threshold (**B**, 1 ms). In the entire recorded population of units, we observed a large spread of gap thresholds ranging over the entire presented range (Figure 3). However, 22.1 % (55 out of 261) of the units showed significant responses to gaps of 1 ms duration already, and only one cell had a significant response at 0.5 ms (Figure 3). Thus, we measured gap thresholds between 0.5 and 1 ms in a notable proportion (>20 %) of units in A1, well in agreement with our behavioral results.

**Figure 2:**
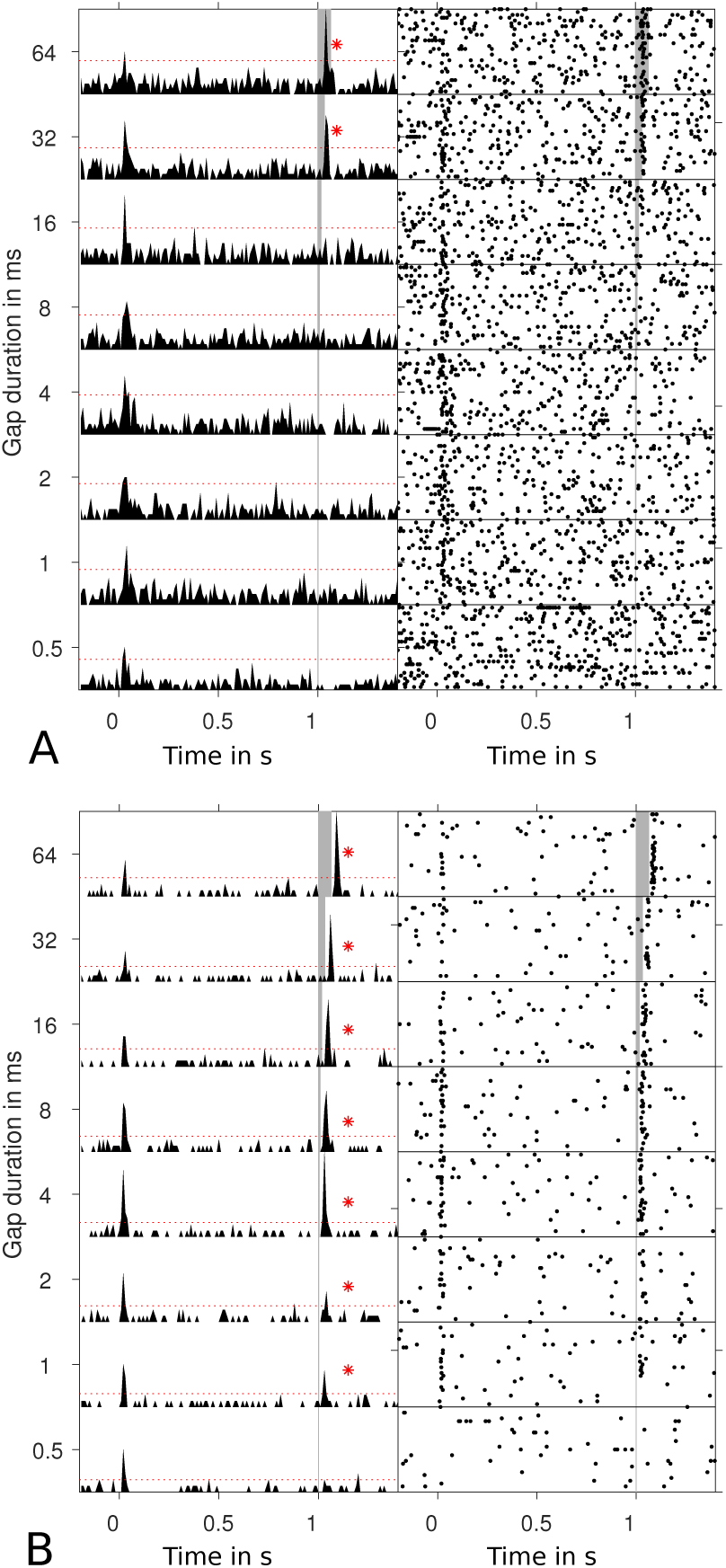
Neural gap detection thresholds in auditory cortex. Single unit examples of neural gap encoding in awake primary auditory cortex. Left hand side depicts peristimulus time histograms of the noise burst starting at t=0 s with a gap of varying duration at t=1 s. Position and duration of the gaps are indicated by the grey areas. Right hand sides show spike-raster plots for the respective units for the same stimuli, number of trials was 30. Red asterisks mark trials with significant responses to the gaps, the shortest gap duration eliciting a response was registered as the threshold (32 ms for the unit in **A** and 1 ms for **B**).

**Figure 3:**
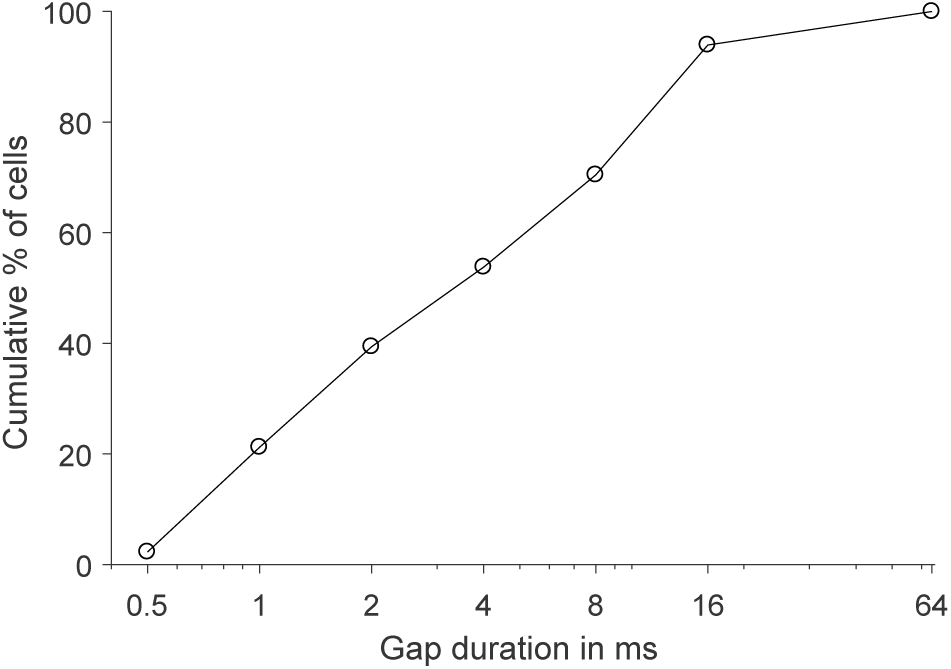
Cumulative distribution of gap thresholds of single units in A1 that responded to any of the gap stimuli (n=261 from 4 animals).

### 3.3 ABR

Since a notable proportion of the cortical population responded with high spiking activity precisely locked to occurrence of gaps of 1 ms and shorter (Figure 2B), we next asked where in the auditory pathway the precise phasic activation after the presentation of a gaps arises from. The information about the presence of a gap is not generated in A1. Either the response pattern of coherently elevated activity is directly inherited from lower stations or it is extracted from the firing pattern that contains information about the gap, for example by a brief interruption of firing during the gap. This point is of central importance for the question of whether gap detection relies on central temporal processing. In order to investigate the representation of GIN at more peripheral stations of auditory pathway, we performed auditory brainstem recordings (ABR). These non-invasive recordings most likely represent highly coherent activity of populations of neurons between different stations. While a peripheral response pattern that is only inherited by A1 should be well observable in the ABR, a more distributed code that has to be interpreted by cortical stations may not be observable in the ABR.

Figure 4 shows ABR curves to GIN stimuli for five different gap durations from a single subject. It can clearly be seen that ABR peaks are temporally related to the gap termination—that is the restart of the noise stream—and not to the start of the gap, which is the noise offset. Therefore, we will draw all further plots of GIN ABRs aligned to the gap termination to facilitate comparisons between ABR curves at different gap durations.

**Figure 4:**
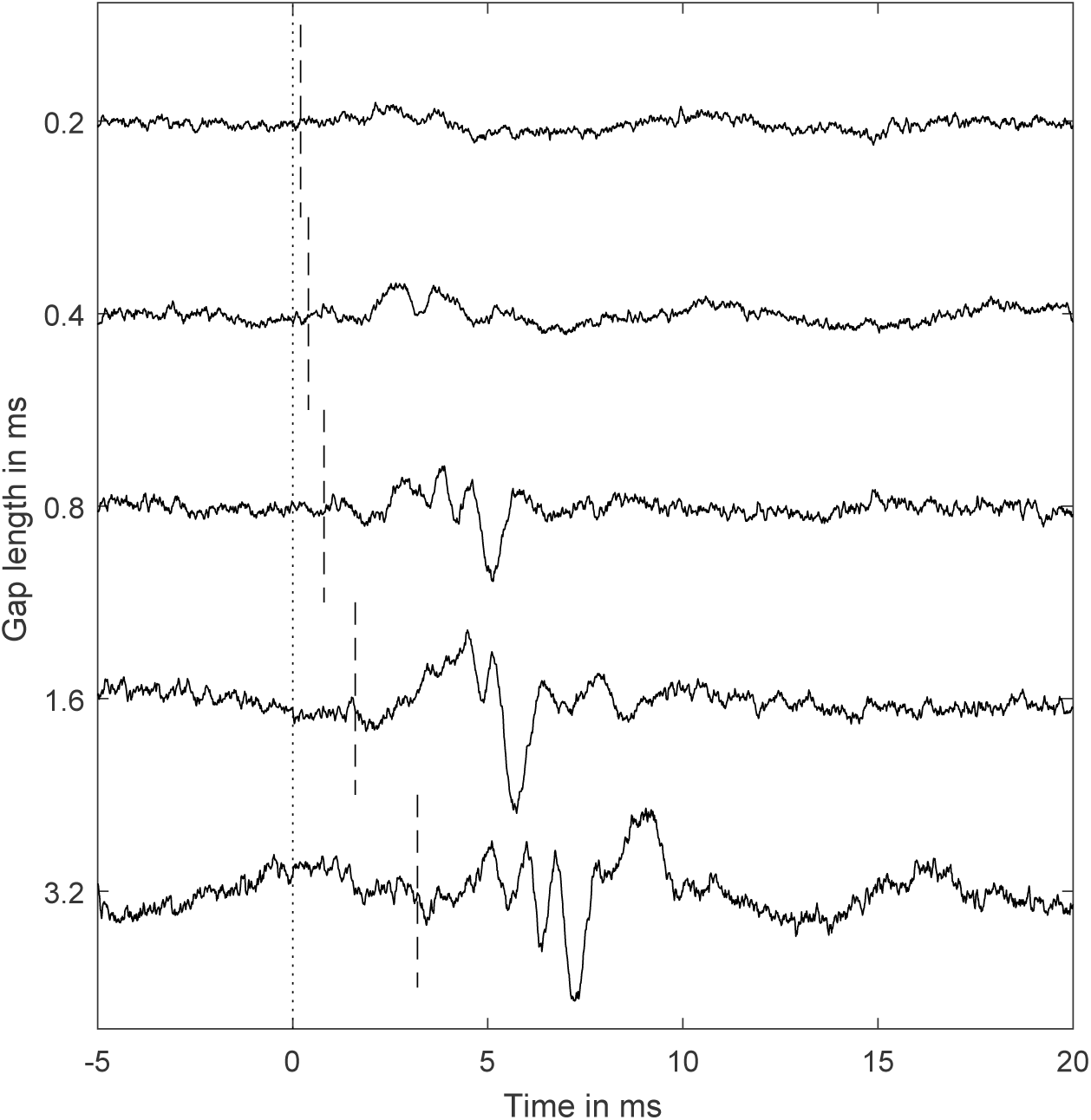
ABRs of a single subject to GIN stimuli for five different gap durations. The dotted line marks the start of the gap (noise offset) and the dashed line marks the gap termination (noise onset). ABR peaks are temporally related to the gap termination and not to its onset.

Figure 5 shows such a plot for a broad range of gap lengths. We enumerated the individual ABR wave components as it was introduced by Jewett and Williston (1971) for the human ABR. Down to a gap duration of around 25 ms, the components I to IV could easily be identified. Until this gap length, the latencies of the components remained stable. For shorter gap durations, all component peak amplitudes decreased and their latencies increased. This affected the individual components to different degrees. The increase of latencies was higher for later than for earlier components. ABR peaks III and IV were least affected by the decrease in amplitude. Therefore, these components are best suited for identifying the presence of a GIN ABR response at short gap lengths.

**Figure 5:**
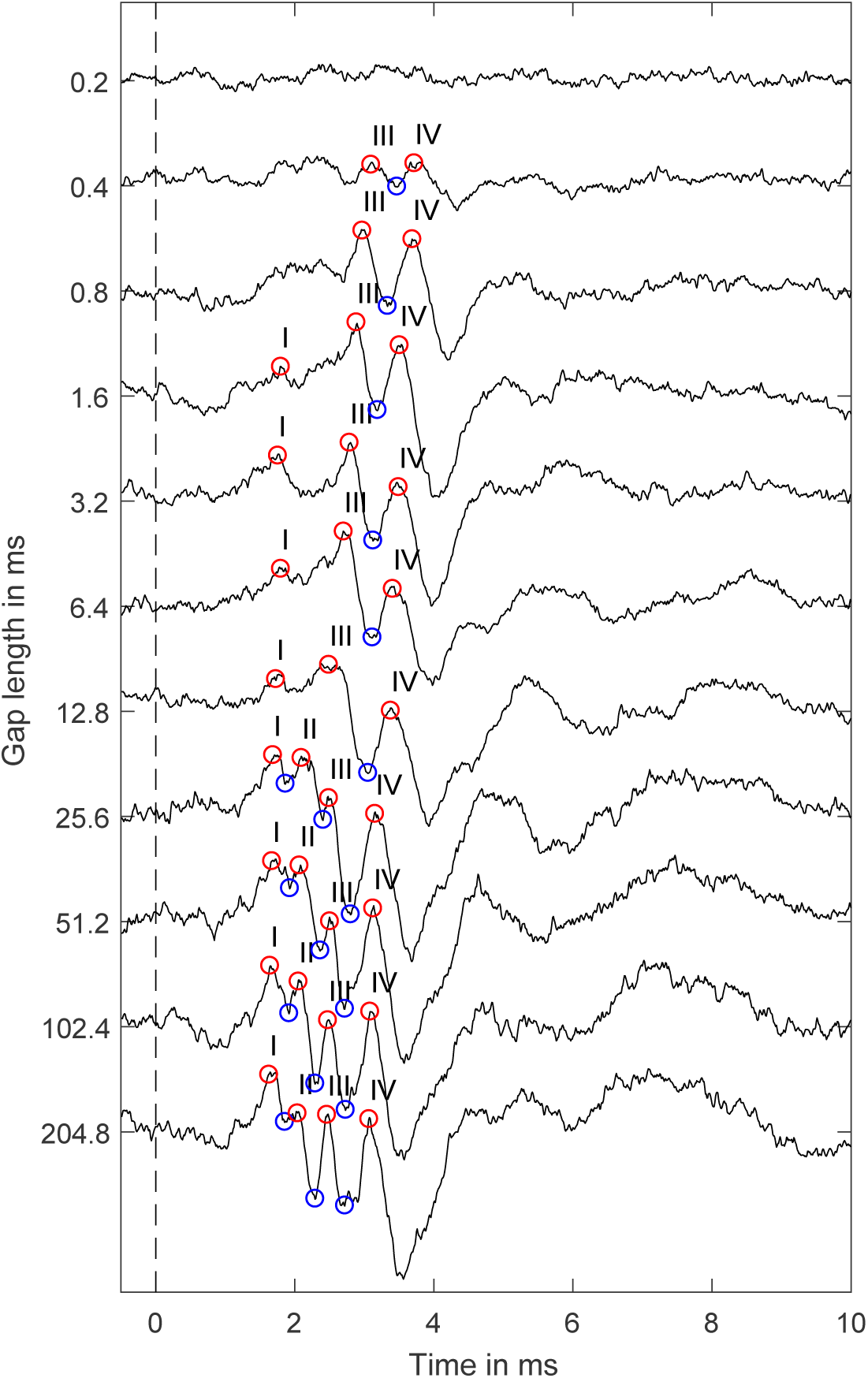
Single subject’s ABRs that follow the gap termination (noise onset, marked with a dashed line) of GIN stimuli. Peaks of the GIN ABRs that might map to those of click/burst evoked ABRs were visually identified (red circles) and labelled according to their click/burst evoked counterparts. With decreasing gap length, the components I and II become increasingly difficult to discern, while the components III and IV remain identifiable down to gap durations of about 0.4 ms. Furthermore, with decreasing gap lengths, peak latencies are increasing.

In order to determine to what extent ABRs to GIN stimuli are comparable to ABRs following NBIS onset responses, we played NBIS stimuli at different levels and recorded the resulting ABR (Figure 6). As expected, long gap durations evoke ABRs that equal noise burst evoked ABRs. Interestingly, the alteration of the ABR curves that we observed with decreasing gap lengths were remarkably similar to those of NBIS at decreasing levels: With decreasing burst levels, peak amplitudes decreased and latencies increased; wave III and IV were the least to vanish.

**Figure 6:**
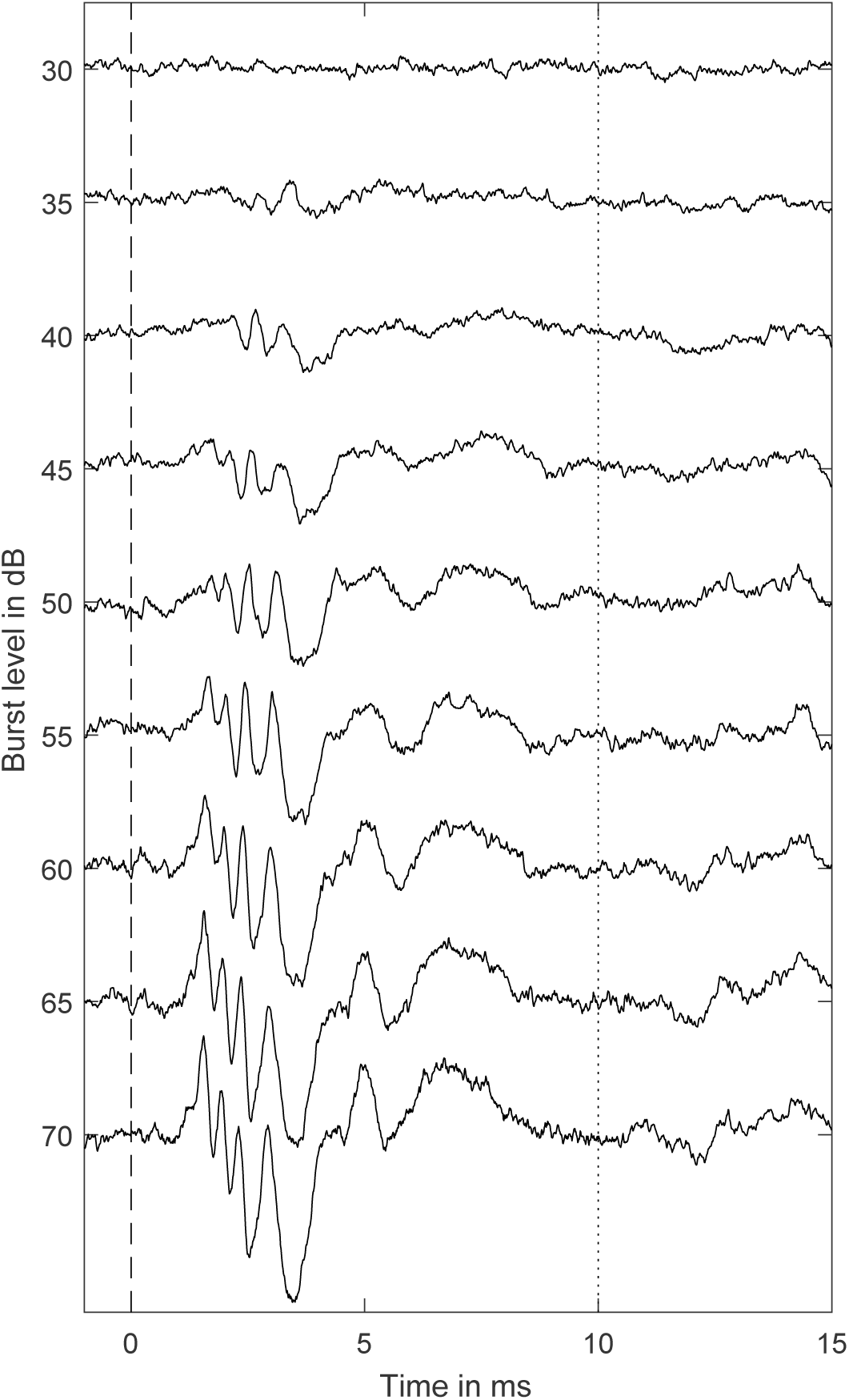
ABRs to NBIS stimuli. The shape of ABRs to NBIS is similar to that of GIN stimuli. Furthermore, the transformation of the ABR’s shape that happens when decreasing the level of NBIS stimuli resembles to that, when decreasing the gap length of GIN stimuli.

In Figure 7, we plotted a noise burst ABR on top of an ABR to a 204.8 ms GIN stimulus. To create this figure, we searched for the NBIS level whose resulting ABR peak latencies best matched those of GIN ABR. Strikingly, this comparably long gap that we played within 60 dB SPL noise evoked a response that was more similar to a 55 dB SPL noise burst than to a 60 dB SPL noise burst. Obviously, the gap length of 204.8 ms was not sufficient to reset the auditory system back into the same state when playing noise bursts in silence. With decreasing gap lengths, the amplitudes of the ABR peak components became lower and their latencies increased.

**Figure 7:**
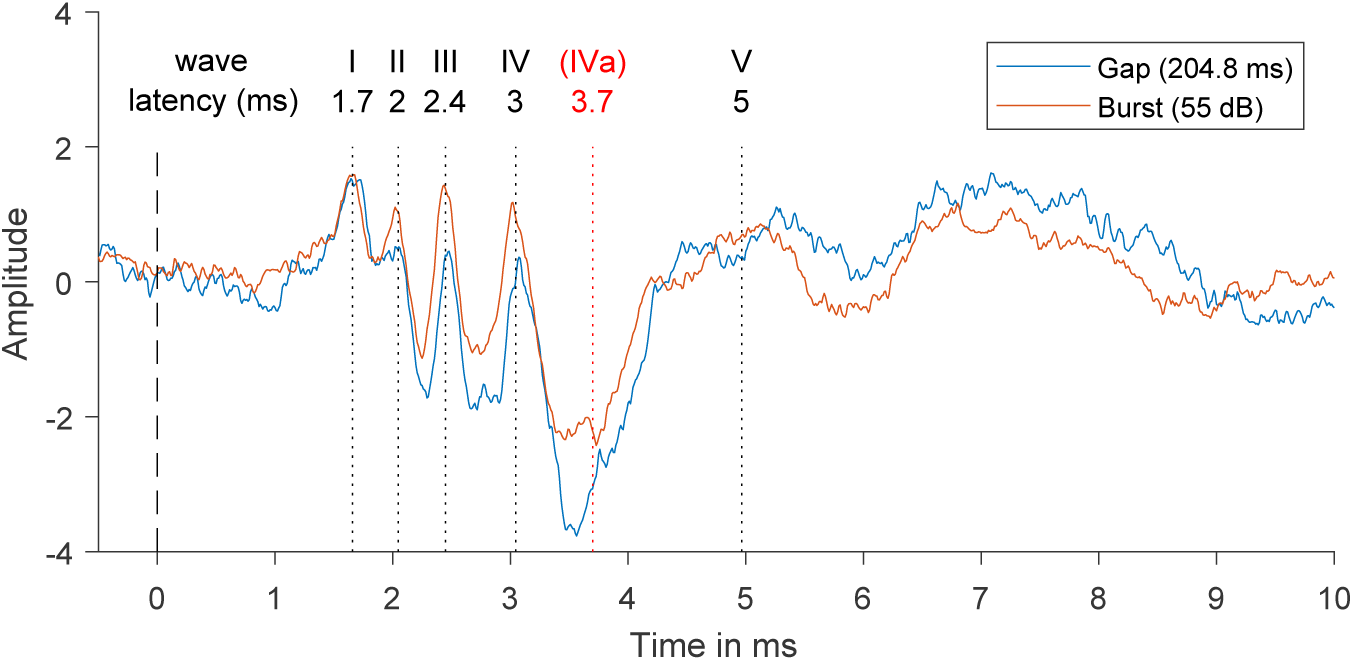
Comparison between a GIN ABR with a long gap duration of 204.8 ms (blue line) and a noise burst evoked ABR (red line). The dashed line marks the noise onset, which is also the gap termination for the GIN stimulus. Dotted lines show visually identified peaks according to the common ABR nomenclature. The GIN ABRs conforms to noise burst evoked ABR. To create this figure, we needed to record ABRs for noise bursts at different levels first, because their peak latencies depend on the level. Here, peak latencies of the ABR evoked from the 204.8 ms gap inserted into 60 dB SPL noise were best mimicked by the ABR that followed a 55 dB SPL noise burst.

Mean ABR curves for GIN stimuli from the same subjects that were used during the behavioral GIN detection experiments, are shown in Figure 8. ABR peaks were standing out from the background activity starting at gap durations of 0.6 ms upwards. Thus, GIN ABR thresholds conform to operant GIN thresholds.

**Figure 8:**
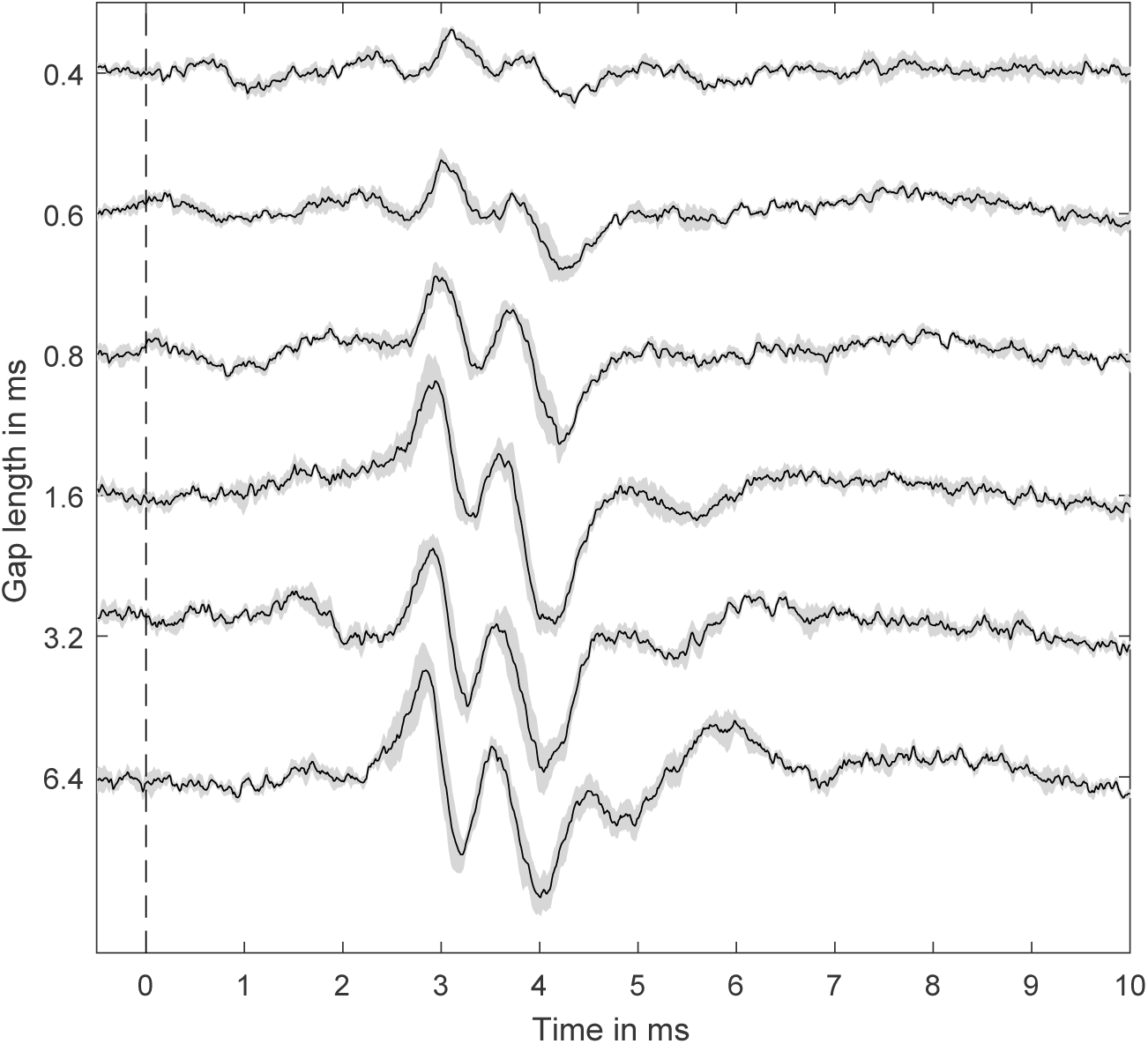
Mean ABR curves for GIN stimuli from the same subjects that were used during the behavioral GIN detection experiment (N=4). The dashed line marks the gap termination (noise onset). The grey area displays ± one SEM. Starting at gap durations of 0.6 ms upwards, ABR peaks stand out from the background activity.

### 3.4 Behavioral performance in aged animals

The measurement of behavioral GIN detection was repeated when the mice were 44 weeks old. We had to exclude two sessions from one animal, one because it exceeded the maximum false alarm rate, and one, because the minimum *d′* value was not reached at any gap duration category. We needed to exclude a single session from another animal, because it exceeded the maximum false alarm rate. We did not observe any age-related degradation of gap detection performance at the follow up measurement. The lines of the mean detection performance across subjects is almost parallel to that of the previous measurement, when the mice were 25 weeks old (Figure 9).

**Figure 9:**
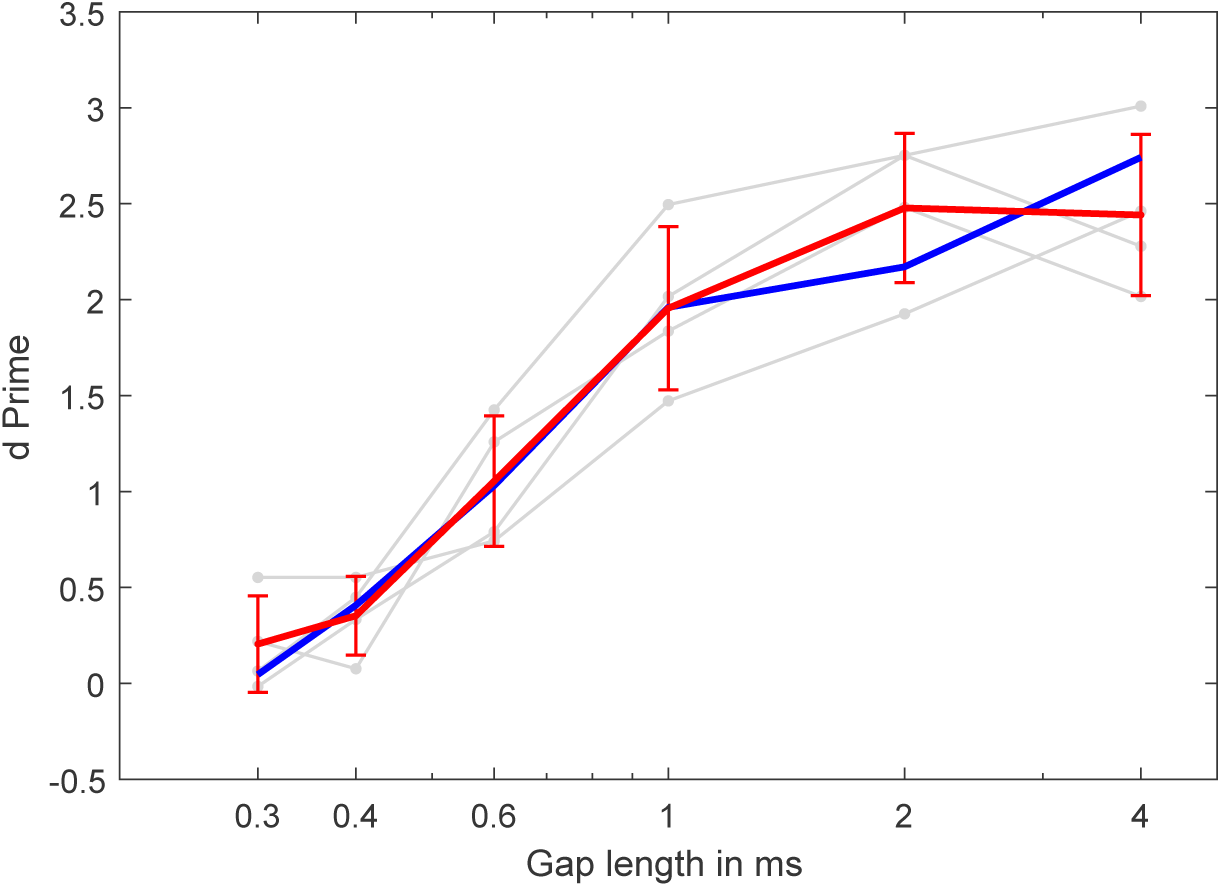
Follow-up measurement of behavioral GIN detection. At the time of the measurement, the mice (N=4) were 44 weeks old. Grey lines show *d′* values for individual subjects. Mean values with error bars representing ± one SEM are painted in red. For comparison, the mean *d′* values of the preceding measurement are included in blue.

## 4 Discussion

We explored gap detection resolution in behavioral, in peripheral electrophysiological (ABR), and in cortical electrophysiological experiments. We found behavioral detection thresholds at gap durations around 0.6 ms (Figure 1). ABR measurements suggest thresholds at comparable gap durations or slightly lower (0.4–0.6 ms, Figure 5 and Figure 8). From the neurons in the AC that respond to GIN, 20 % responded to gap durations of 1 ms and lower (Figure 3). Behavioral gap detection resolution did not change with age (Figure 9).

### 4.1 Comparison to gap detection thresholds reported by previous animal studies

In comparison to all other behavioral GIN detection experiments with mice that we are aware of, our mice had superior detection thresholds (Walton, Frisina, Ison, and O’Neill 1997; Ison, Allen, Rivoli, and J. T. Moore 2005; Radziwon et al. 2009). We can think of three reasons for that: First, most studies employ the technique of prepulse inhibition (PPI), which is known to be less sensitive than operant conditioning (Behrens and Klump 2015; Lauer, Behrens, and Klump 2017). Second, we embedded our gaps in noise that had a broad spectrum. Therefore, our noise possibly had more energy within the higher hearing range of the mice, which is most important for gap detection (Ison, Allen, Rivoli, and J. T. Moore 2005). Third, B6.CAST mice are among the mice with the best hearing, with lower threshold than commonly used CBA mice (Zheng, Johnson, and Erway 1999).

### 4.2 Similarity of responses of brief GINs and noise bursts

Our recordings of ABRs show that the response to a GIN can be regarded as a response to the onset of the noise after the termination of the gap. The transformation of the ABR curve when the gap length was decreased was similar to that of NBIS when the burst level was decreased indicating a smaller coherently firing population. We suggest that a similar process underlies the ABR to GIN. With decreasing gap length, the population of neurons that responds to the post-gap onset decreases. The similarity of encoding post-gap onsets and noise bursts hast already been reported at different levels of the auditory pathway (Guo and Burkard 2002; Keller, Kaylegian, and Wehr 2018).

Our data suggest that the detection of GIN does not require complex neural computation. A simple and strong neural representation of the gap stimulus exists already at peripheral and early central stages of the auditory pathway. Other studies suggest the central stations (AC) to play a crucial role in behavioral GIN detection (Syka, Rybalko, Mazelová, and Druga 2002; Ison, O’Connor, Bowen, and Bocirnea 1991; Weible, Liu, Niell, and Wehr 2014). Especially loss of functional inhibition at central stages has been linked to impaired gap detection in aged animals (Gleich, Hamann, Klump, Kittel, and Strutz 2003). Since cortical inhibition is functionally diminished in aged mice (Martin del Campo, Measor, and Razak 2012), this decrease may affect cortical gap representation as well. Indeed, a subpopulation of cortical inhibitory units has recently been found to be especially strongly modulated by gaps (Keller, Kaylegian, and Wehr 2018). However, this modulation was not specific to GINs but also for onset of noise bursts.Boosting of inhibition may nonspecifically increase signal-to-noise ratios at central stages (Natan et al. 2015), thereby improving detectability of brief gaps (Gleich, Hamann, Klump, Kittel, and Strutz 2003) but also similar non-salient targets. Conversely, an age-related loss of central inhibition may hamper low signal-to-noise signals, including, but not exclusive for short GINs.

### 4.3 Aging and temporal processing deficits

In the group of mice that we measured twice during their life span, GIN detection performance did not decrease with age. Besides, their ABR to GIN stimuli at middle age is indistinguishable from that of younger individuals. This finding contrasts with most studies that investigated the effect of aging on gap detection. One reason for the disagreement of our data and previous studies may be that we used middle-aged mice (15 month old) while other studies used older animals (e.g. Walton, Frisina, and O’Neill 1998). The reason for our choice was to ensure good peripheral hearing at an age for which age-related changes both in the encoding of gaps (Williamson, Zhu, Walton, and Frisina 2015) and central physiology (Martin del Campo, Measor, and Razak 2012) had been reported before. It may thus be possible that the CAST line used here develops gap-detection deficits at older age, but this likely goes along with peripheral hearing loss as well.

Studies that investigate auditory temporal processing deficits at central sites in aged humans generally face the problem that sensorineural hearing loss for high frequency regions is widespread (Agrawal, Platz, and Niparko 2008). Gap detection performance in mice also strongly depends on hearing in the high-frequency range (Ison, Allen, Rivoli, and J. T. Moore 2005). Thus, high-frequency hearing loss might be a confounding factor in animal experiments as well.

Our data on middle aged mice does not contradict a scenario, in which peripheral and central consequences of aging jointly cause a loss in temporal resolution. Central mechanisms may partly be able to compensate a loss in temporal precision of upstream inputs (Walton, Frisina, and O’Neill 1998; Gleich, Hamann, Klump, Kittel, and Strutz 2003; Kopp-Scheinpflug et al. 2011; Anderson and Linden 2016; Keller, Kaylegian, and Wehr 2018. The most popular mouse strain used in auditory research, CBA mice, is commonly regarded as not developing age-related peripheral hearing loss (Zheng, Johnson, and Erway 1999). However, Spongr (1997) found significant loss of significant amounts of outer and inner hair cells in CB mice. This loss may be partly responsible for the difference between the present study and others using CBA mice (Williamson, Zhu, Walton, and Frisina 2015). We suggest that a hampered peripheral encoding of gap-offsets in many animal models may reduce the neural representation of short GINs in many experimental models of aging. Central mechanisms that enhance neural representation of low signal-to-noise signals including, but not limited to GINs, may then play a major role to determine perceptual salience.

Here, we found highly precise gap-end encoding in the periphery at an age where gap-encoding deficits have been found in other mouse strains (Williamson, Zhu, Walton, and Frisina 2015). Our conclusion is that central deficits alone may not primarily be responsible for deficits in gap detection, at least not at middle age.

### 4.4 Gap detection and hidden hearing loss

One of the main factors for further processing of short GIN maybe their precise encoding in the periphery. The most common method to compensate for sensorineural hearing loss as a confounding factor during the investigation of central presbycusis is to use sound stimuli that are well above audiometrically derived hearing thresholds. In the presence of hidden hearing loss this compensation may not be effective. Hidden hearing loss is noise-induced or age-related cochlear neuropathy in the absence of hair cell loss (Viana et al. 2015) and is believed to be contributing to impaired speech intelligibility and other listening situations that involve temporal auditory processing (Liberman, Epstein, Cleveland, Wang, and Maison 2016; Plack, Barker, and Prendergast 2014). Most notably, CBA mice develop cochlear synaptic loss that progresses with age long before their number of hair cells decreases (Sergeyenko, Lall, Liberman, and Kujawa 2013), which may explained increased ABR thresholds at middle age already (Williamson, Zhu, Walton, and Frisina 2015). Up to now, temporal processing deficits in aged subjects with normal audiograms was commonly attributed to central processing deficits. Hidden hearing loss, which is a peripheral disorder, could be an alternative explanation for temporal processing deficits in aged human, CBA mice, and certainly other species with normal audiograms.

### 4.5 Conclusion

Age-related sensorineural hearing loss is a confounding factor in experiments that investigate temporal processing deficits in central presbycusis. Solely accounting for differences in audiograms between younger and older subjects to minimize the influence of this factor is insufficient. CAST mice, a mouse line that does not develop age-related sensorineural hearing loss, provide an alternative.

We found ultra-fine auditory gap resolution in this mouse line. The gap resolution is preserved at middle age, suggesting that gap detection is primarily determined by peripheral, not central processing.

The neural representation of GINs is an excitatory response on the onset of the noise that follows the gap termination. This response can be observed at peripheral, early, and late central (AC) stages along the auditory pathway, without being strongly transformed at any stage. The response is comparable to that of a NBIS, with decreasing gap lengths being comparable to decreasing intensities in NBIS stimuli. The rather simple representation of GIN does not involve complex neural computation, which makes it unlikely that later auditory stages are critical in determining GIN resolution.

Instead, we propose that previous findings indicating impaired gap representation at central stages result either form decreased peripheral sensitivity to high frequencies or from central deficits that hamper the detection of low signal-to-noise signals in general.

Representation and detection of GIN may thus not be the ideal single indicator of central temporal processing.

## Acknowledgments

We thank G. Klump for comments on a previous version of this manuscript, I. Rauser for conducting behavioral experiments, and R. Beutelmann for providing DSP software components that were used in our ABR measurement system.

